# Toll-like receptor 3 activation increases voluntary alcohol intake in C57BL/6J male mice

**DOI:** 10.1101/476457

**Authors:** Anna S. Warden, Moatasem M. Azzam, Adriana DaCosta, Sonia Mason, Yuri A. Blednov, Robert O. Messing, R. Dayne Mayfield, R. Adron Harris

## Abstract

Many genes differentially expressed in brain tissue from human alcoholics and animals that have consumed large amounts of alcohol are components of the innate immune toll-like receptor (TLR) pathway. TLRs initiate inflammatory responses via two branches: (1) MyD88-dependent or (2) TRIF-dependent. All TLRs signal through MyD88 except TLR3. Prior work demonstrated a direct role for MyD88-dependent signaling in regulation of alcohol consumption. However, the role of TLR3 as a potential regulator of excessive alcohol drinking has not previously been investigated. To test the possibility TLR3 activation regulates alcohol consumption, we injected mice with the TLR3 agonist polyinosinic:polycytidylic acid (poly(I:C)) and tested alcohol consumption in an every-other-day two-bottle choice test. Poly(I:C) produced a persistent increase in alcohol intake that developed over several days. Repeated poly(I:C) and ethanol exposure altered innate immune transcript abundance; increased levels of TRIF-dependent pathway components correlated with increased alcohol consumption. Administration of poly(I:C) before exposure to alcohol did not alter alcohol intake, suggesting that poly(I:C) and ethanol must be present together to change drinking behavior. To determine which branch of TLR signaling mediates poly(I:C)-induced changes in drinking behavior, we tested either mice lacking MyD88 or mice administered a TLR3/dsRNA complex inhibitor. MyD88 null mutants showed poly(I:C)-induced increases in alcohol intake. In contrast, mice pretreated with a TLR3/dsRNA complex inhibitor reduced their alcohol intake, suggesting poly(I:C)-induced escalations in alcohol intake function are, at least partially, dependent on TLR3. Together, these results strongly suggest that TLR3-dependent signaling drives excessive alcohol drinking behavior.

**Highlights:**

- Activation of TLR3 *via* poly(I:C) increased alcohol intake.
- Poly(I:C) and ethanol must be present together to change drinking behavior.
- Increased alcohol intake due to poly(I:C) is independent of MYD88.
- Increased alcohol intake due to poly(I:C) is dependent on TLR3.

## 1. Introduction

Alcohol consumption activates peripheral and central inflammatory pathways leading to increases in innate immune signaling (1–3). Inflammatory and immune-related signaling genes are differentially expressed in the frontal cortex of human alcoholics (4, 5), selectively-bred high drinking mice (6) and ethanol-exposed mice (7–10), indicating a role for innate immune signaling in regulating alcohol intake. Moreover, proinflammatory cytokines have been detected in the serum of human alcoholics and higher levels of cytokines correlated with alcohol craving (11). Taken together, this suggests that inflammatory and immune-related signaling may regulate alcohol craving and consumption. Behavioral validation studies confirmed that mice with null mutations in different immune-related genes drink less ethanol, whereas activation of innate immune signaling using lipopolysaccharide (LPS) increases ethanol intake (12, 13). Many of the innate immune genes implicated in these studies mediate their effects through activation of toll-like receptors (TLRs).

TLRs recognize a broad range of molecular motifs known as pattern-associated molecular patterns (PAMPs) to initiate innate immune responses (14). TLRs initiate inflammatory responses via two intracellular signaling transduction cascades: (1) Myeloid differentiation response gene 88 (MyD88)-dependent or (2) TIR-domain-containing adapter-inducing interferon β (TRIF)-dependent. Only TLR3 initiates inflammatory responses solely through the TRIF-dependent pathway.

Prior studies indicate a direct role for TLR signaling in the regulation of alcohol intake. Specifically, the field has focused on TLR4/MyD88-dependent signaling (12, 15–19). For instance, activation of TLR4 via LPS increased alcohol intake; whereas, siRNA inhibition of the downstream MyD88-dependent signaling component IKKβ decreased alcohol consumption (12, 19). However, despite evidence that TLR4/MyD88-dependent signaling is important for alcohol responses, it was recently found that TLR4 is not a critical determinant of excessive drinking in animal models (20, 21). Moreover, a recent study could not replicate that LPS pretreatment increases alcohol consumption in continuous two-bottle-choice or drinking-in-the-dark (22). This suggests that other innate immune pathways may be important in the regulation of excessive alcohol consumption. We hypothesize that TLR3/TRIF-dependent signaling may be a key determinant of excessive drinking.

Increased expression of TRIF-dependent pathway transcripts and proteins has been found in the frontal cortex and nucleus accumbens of male mice subjected to chronic voluntary alcohol consumption—the largest increases in TRIF-dependent pathway components were in the medial prefrontal cortex, which is why we chose this brain region for subsequent analysis (23). Increased expression of TRIF-dependent components has also been found in postmortem samples of cerebral cortex and basolateral amygdala from human alcoholics (24–26). TLR3 transcript abundance correlates with lifetime alcohol consumption in human alcoholics (25). Chronic ethanol exposure combined with TLR3 stimulation increases expression of proinflammatory cytokines in mouse frontal cortex (27). Inhibiting the downstream TRIF-signaling components IKKI and TBK1 reduces ethanol consumption in male mice, suggesting that an intact TRIF-dependent pathway in necessary for alcohol consumption (24). However, it is not known if activating TRIF-dependent signaling increases excessive drinking behavior.

In this study, we tested the hypothesis that activation of TLR3-dependent signaling increases alcohol intake. We activated TLR3-dependent signaling by administering the TLR3 agonist polyinosinic-polycytidylic acid (poly(I:C)) to C57BL/6J mice. We found that poly(I:C) increased ethanol intake through an interaction with ethanol that is, at least partially, dependent on TLR3. We also documented changes in innate immune transcripts after chronic poly(I:C) administration during chronic alcohol intake that may mediate increased alcohol consumption. These results establish TLR3-dependent signaling as an important regulator of alcohol consumption in male mice. Inhibiting TLR3-dependent signaling may be a novel strategy to reduce excessive alcohol drinking and the brain immune response to chronic ethanol exposure.

## 2. Materials and Methods

### 2.1 Mice

Generation of *Myd88* (B6.129P2(SJL)-*Myd88tm1.1Defr*/J, stock #009088) knockout (KO) mice was performed as described previously (28). Mutant strain purchased from Jackson Laboratories was backcrossed on a C57BL/6J genetic background more than 6 generations. Male C57BL/6J mice were originally purchased from Jackson Laboratories, Bar Harbor, ME at 8-10 weeks of age and bred to maintain our colony. Behavioral testing began when the mice were at least 2 months old, and mice were weighed every 4 days. All experiments were conducted in isolated behavioral testing rooms. The University of Texas at Austin Institutional Animal Care and Use Committee approved all experiments.

### 2.2 Drug administration

Poly(I:C) HMW obtained from Invivogen (San Diego, CA) was prepared as previously described (29). To determine the time course of poly(I:C), we injected poly(I:C) (5mg/kg) intraperitoneally (i.p.) and then sacrificed mice at 3, 24, and 48 hours post-injection before qRT-PCR analysis. The 5mg/kg dose was chosen to minimize the sickness response (29) and mitigate any adverse effect on voluntary ethanol consumption 24-36 h post-poly(I:C) injection. For ethanol drinking studies, poly(I:C) (2, 5, or 10mg/kg, i.p.) was administered every four days during no alcohol access periods. For the control injection, 0.9% saline (volume matched) was administered to control groups. Single use, sterile needles (27.5 gauge) were used. All injections we made between 8 am and 9 am to animals 8-16 weeks of age.

TLR3/dsRNA complex inhibitor obtained from Sigma Aldrich (St. Louis, MO) was prepared for injection in 10% DMSO as previously described (30, 31). For qRT-PCR experiments, we injected 2mg inhibitor, waited 30 minutes and then injected poly(I:C) (5mg/kg, i.p.) and sacrificed mice at 3 hours post-injection. To test the effect of the inhibitor on poly(I:C)-induced escalations in alcohol consumption, mice were injected with poly(I:C) every four days, and underwent two-bottle choice every-other-day (EOD) drinking for a total of 36 days. After we verified that poly(I:C) increased alcohol intake, we injected 2mg of inhibitor, waited 30 minutes and then injected poly(I:C) (5mg/kg, i.p.) and allowed animals to undergo eight days EOD drinking with injections every four days. To test the effect of the inhibitor alone on alcohol consumption, the 2mg of inhibitor or saline (10% DMSO) was injected every four days for a total of 24 days during EOD drinking. To control for DMSO drug dilutions, saline (10% DMSO, volume matched) was administered to both the saline/saline and the saline/poly(I:C) groups.

### 2.3 Two-bottle choice every-other-day procedure

Intermittent EOD access to ethanol increases voluntary drinking in rats (32, 33) and mice (34–36). Mice were given EOD access to ethanol (15 or 20%) as previously described (12). The quantity of ethanol consumed was calculated as g/kg body weight/24 h. Total fluid intake was calculated as g/kg body weight/24h. For all EOD access drinking experiments, each group had ten animals, (n=10/group).

### 2.4 Preference for saccharin

Mice were tested for saccharin consumption using an EOD protocol in which one bottle contained water and the other contained saccharin solution. Mice were offered increasing concentrations of saccharin (0.0008 and 0.0016%). Each concentration was offered for a series of four poly(I:C) injections (16 days). Bottle positions were changed for each saccharin session. (n=10/group).

### 2.5 Brain and organ collection

For qRT-PCR experiments, brains were quickly harvested and the prefrontal cortex rapidly dissected before being snap frozen in liquid nitrogen. For immunohistochemistry experiments, mice were anesthetized with isoflourane, transcardially perfused with 0.9% saline until cleared of blood, and then perfused with freshly prepared 4% paraformaldehyde PFA in phosphate-buffered saline (PBS). Immediately thereafter, the brain and liver were removed and post fixed in 4% (PFA) at 4□°C for 24□h followed by cryoprotection for 24□h at 4□°C in 20% sucrose. Brains and livers were then placed in plastic molds containing optimum cutting temperature compound (OCT, VWR, Radnor, PA) and quickly frozen in isopentane on dry ice.

### 2.6 qRT-PCR

Total RNA was isolated, quantified and converted to cDNA as described (9). Relative quantification of mRNA levels was determined using BIORAD software as previously described (9, 10, 37). *Gusb* was selected as an endogenous control to normalize target gene mRNA levels. For all qRT-PCR, six animals were used per group, unless otherwise noted.

### 2.7 Immunohistochemistry

Thirty-micron sections from brain or liver were treated for immunohistochemistry, as described (37). Sections incubated with K1 (mouse α dsRNA, 1:200, Scicons, Hungary) and Alexa Fluor 488 (donkey α mouse, 1:1000, A21202, Invitrogen, Carlsbad, CA). Sections were mounted in 0.2% gelatin, dehydrated, and cover slipped with a DAPI (4’,6-diamidino-2-phenylindole)-containing mounting medium (Vector Labs, Burlingame, CA). For all immunohistochemistry experiments, four animals were used per group, unless otherwise noted.

### 2.8 Microscopy and quantitative image analysis

Identification of dsRNA positive cells was performed using a Zeiss Axiovert 200□M fluorescent light microscope for both brain and liver (Zeiss, Thornwood, NY) equipped with an Axiocam b/w camera as previously described for prefrontal cortex (37). Parameters used for image acquisition were identical across treatments.

To quantify K1 synthetic dsRNA counts and fluorescence intensity in prefrontal cortex and liver, a single in-focus plane was acquired and analyzed as previously described using Image J (v1.49u, NIH) (37, 38).

### 2.9 Antibody controls

We tested the specificity of the primary antibody by injecting animals with a ssRNA agonist (R848, 200nmol, Invivogen, San Diego, CA), which should have no signal when using the dsRNA K1 antibody (Supplemental Figure 1A). We also tested the specificity of the secondary antibody (secondary control) by replacement of the primary antibody with only the serum of the appropriate species (Supplemental Figure 1B) or by replacement of both primary and secondary antibody with serum of the appropriate species (Supplemental Figure 1C). Furthermore, K1 has been widely used and its specificity has been previously validated (39–41).

**Figure 1:**
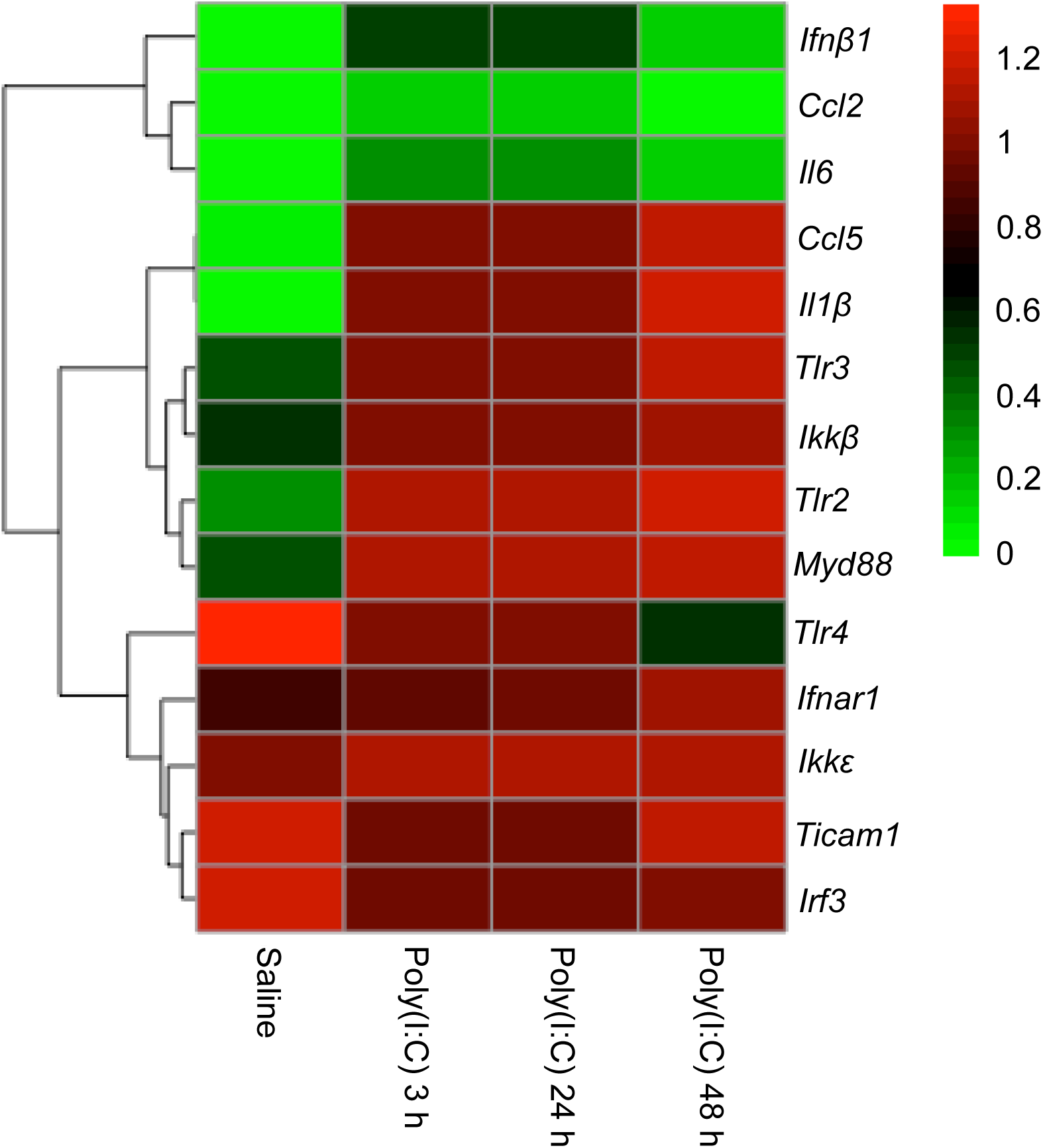
Poly(I:C) increases innate immune mRNA levels in prefrontal cortex. Heat map showing transcript abundance at four time points, before and following administration of poly(I:C). The transcript levels are presented using fold-change values (Log2 format) normalized to endogenous control. The red and green colors indicate high and low transcript abundance, respectively. The scale representing the relative signal intensity values is shown on the right. Data were analyzed by hierarchical clustering, n=6 per group.

### 2.10 Statistical analysis

Data are reported as mean ± SEM values, unless otherwise noted. The statistics software program GraphPad Prism (GraphPad Software, Inc., La Jolla, CA) was used to perform 2-way ANOVAs, Pearson correlations and Student's *t*-tests. Drinking data were analyzed by repeated-measures 2-way ANOVA followed by Bonferroni post-hoc tests. Transcript abundance data were analyzed by two-way ANOVA followed by Tukey’s HSD post-hoc tests. The Pearson correlation (α = 0.05) was used to evaluate correlations between ethanol consumption and transcript abundance. Grubbs test (α = 0.05) was used to detect potential outliers. Student’s t-tests (two-tailed) were used to analyze raw qRT-PCR data, dsRNA immunopositive cell counts, and the TLR3/dsRNA inhibitor drinking experiment.

## 3. Results

### 3.1 Poly(I:C) rapidly increases innate immune transcript abundance in the prefrontal cortex

Poly(I:C) induces a pro-inflammatory response in the central nervous system (27, 29, 42). We hypothesized that this proinflammatory response is due at least in part to increased TRIF-dependent pathway signaling, which would be reflected in increased abundance of pathway transcripts and transcriptional outputs. To capture changes over time, we measured the time course of poly(I:C) effects on prefrontal cortical transcripts for TRIF- and MyD88-dependent pathway members and for select cytokines and chemokines implicated in behavioral responses to ethanol (see (43) for review). Poly(I:C) (5mg/kg, i.p.) produced three distinct patterns of change in transcript levels: (1) genes dynamically changed by poly(I:C) across time, (2) genes persistently increased across time, and (3) genes unchanged by poly(I:C) (Figure 1, Supplemental Figure 2). Transcripts for the proinflammatory mediators *Ifn*β*1, Il1*β*, Ccl2,* and *Il6* showed a strong increase that peaked at 3 hours and returned to baseline by 48 hours [F_treatment × time_ (2,30) = 18.52 (p<0.0001) for *Ifn*β*1*, 5.54 (p<0.001) for *Il1*β, 14.59, (p<0.0001) for *Ccl2*, and 14.58 (p<0.0001) for *Il6*]. In contrast, the transcript for *Tlr4* was unique in that it decreased at 48 hours after poly(I:C) treatment [F_treatment × time_(2,30)=3.89, p=0.03]. Transcripts for *Tlr3, Tlr2, Myd88*, *Ikk*β*, and Ccl5* followed a different course and were elevated at all time points [F_treatment_(1,30)=108.7, (p<0.0001) for *Tlr3*, 70.8 (p<0.0001) for *Tlr2*, 44.07 (p<0.0001) for *Myd88*, 30.04 (p<0.0001) for *Ikk*β and 242.3 (p<0.0001) for *Ccl5*]. In contrast to the downstream TRIF pathway member *Ccl5*, other TRIF-dependent pathway components (*Ticam1, Ikk*ε*, and Irf3*) were unchanged by poly(I:C) treatment. These findings suggest that activation of TLR3 signaling not only increases levels of TRIF-dependent pathway outputs *Ifn*β*1* and *Ccl5*, but also of several innate immune genes that function outside of the TRIF-dependent pathway.

Poly(I:C) is a long dsRNA that can be recognized by either TLR3 or MDA5 (44-45). To determine whether the proinflammatory response after poly(I:C) was dependent on TLR3, we pretreated animals with a TLR3/dsRNA complex inhibitor (2mg, i.p.) followed by poly(I:C) (5mg/kg, i.p.) and measured prefrontal cortical transcripts for select inflammatory mediators implicated in TLR3 response to poly(I:C). TLR3/dsRNA complex inhibition reduced poly(I:C)-induced increases in *Ifn*β*1*, *Ccl5*, *Ccl2*, *Il6*, and *Il1*β (Supplemental Figure 3). This suggests that the inflammatory response after poly(I:C) is, at least partially, mediated by TLR3.

**Figure 2:**
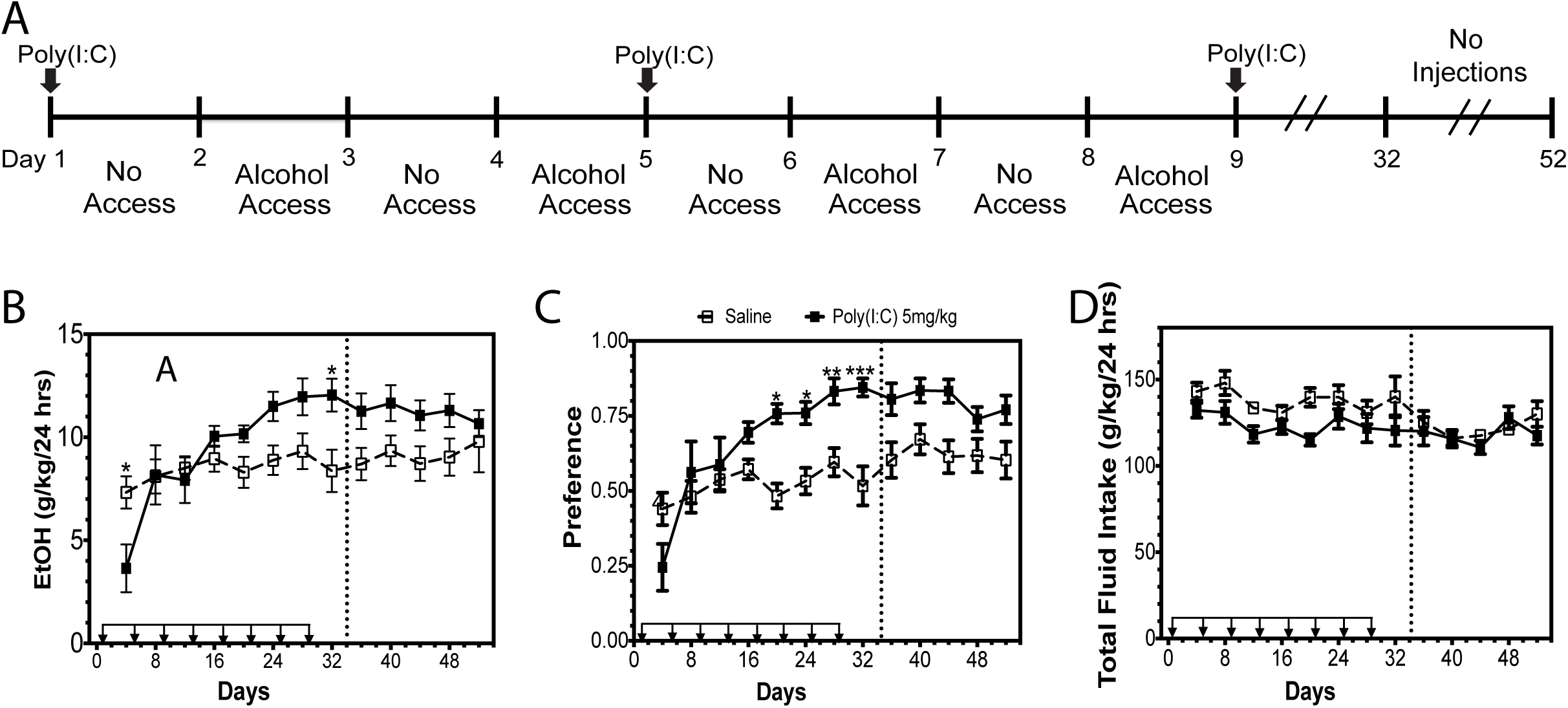
Poly(I:C) increases alcohol intake over time. (A) C57BL/6J male mice were injected with saline or poly(I:C) (5mg/kg) every four days for a total of eight injections during two-bottle choice, every-other-day drinking (EOD,15%). (B) Ethanol (EtOH) intake (g/kg/24 h); (C) preference for EtOH; (D) total fluid intake. Arrows indicate days when animals received injections. Dashed line represents a “no injection” period during which animals continued EOD without injections. Data represented as mean ± s.e.m. (***p<0.0002, **p<0.0021, *p<0.05 for Bonferroni post-hoc tests, n=10 per group).

**Figure 3:**
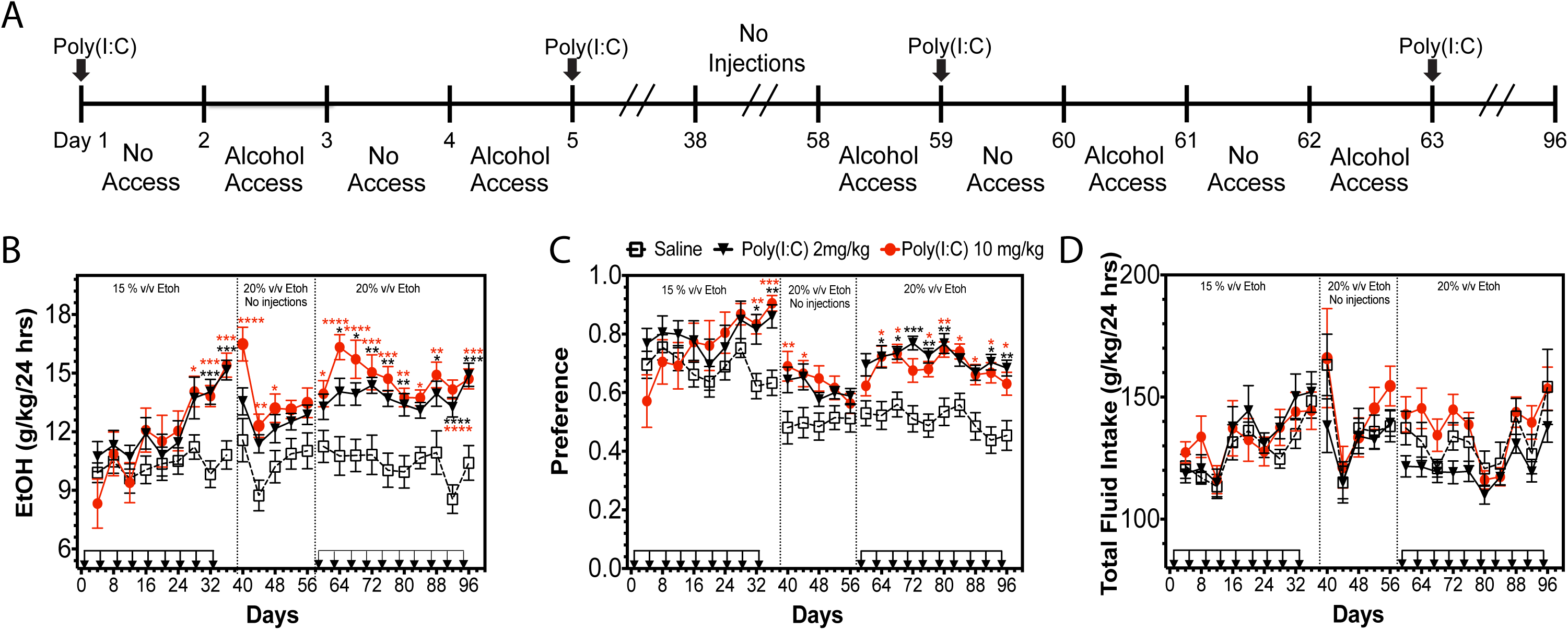
Poly(I:C) increases alcohol intake across multiple doses. (A) C57BL/6J male mice were injected with saline, poly(I:C) (2mg/kg), or poly(I:C) (10mg/kg) every four days during a two-bottle choice every-other-day drinking procedure (EOD, 15% or 20%). (B) Ethanol (EtOH) intake (g/kg/24 h); (C) preference for EtOH; (D) total fluid intake. Dashed line represents a “no injection” period during which animals continued EOD without injections. Data represented as mean ± s.e.m. (****p<0.0001, ***p<0.0002, **p<0.0021, *p<0.05 for Bonferroni post-hoc tests, n=10 per group).

Poly(I:C) is a polymer, therefore it might not reach the brain when administered by intraperitoneal injection. To investigate whether poly(I:C) is present in brain after systemic administration, we measured its presence in prefrontal cortex and liver using the K1 IgG2a monoclonal antibody, which detects synthetic dsRNA (39–41). Following administration of poly(I:C) (5mg/kg, i.p.), dsRNA-like immunoreactivity was detected in both prefrontal cortex and liver (Supplemental Figure 4). This finding suggests poly(I:C) does reach the brain after intraperitoneal administration, and direct activation of TLR3 in the central nervous system contributes to its effects.

### 3.2 Poly(I:C) increases alcohol intake over time

Poly(I:C) can increase abundance of innate immune transcripts in brains of male mice, and other activators of innate immunity like LPS increase alcohol drinking (12), therefore we next tested the hypothesis that TLR3 activation increases alcohol intake. C57BL/6J male mice were administered poly(I:C) (5mg/kg, i.p.) every four days during an intermittent (EOD) consumption procedure (15% ethanol) (Figure 2A). Following the first injection of poly(I:C), there was a transient decrease in ethanol consumption, but after multiple injections, alcohol consumption increased [Figure 2B, F_treatment × time_ (12,216) = 3.68, p<0.0001]. A similar pattern was observed for ethanol preference [Figure 2C, F_treatment × time_ (12, 216) = 4.77, p<0.0001]. Total fluid intake did not change during the course of the experiment [Figure 2D, p=0.26; F_treatment_(1,18)=4.22, p=0.055; F_time_(12, 216)=4.05, p<0.0001; F_treatment × time_(12,216)=1.229].

**Figure 4:**
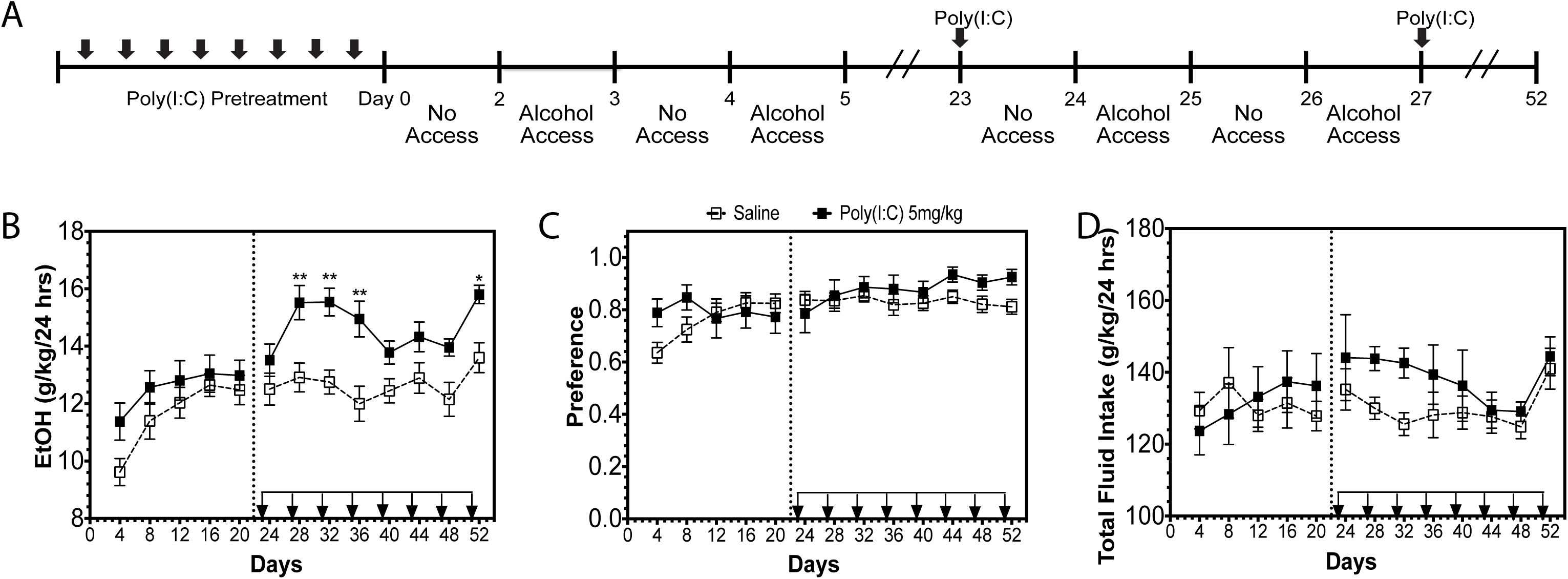
Pretreatment with poly(I:C) does not alter subsequent alcohol intake. (A) C57BL/6J male mice were pretreated with saline or poly(I:C) (5mg/kg) every four days for a total of eight injections before beginning a two-bottle-choice every-other-day procedure (EOD,15%). (B) Ethanol (EtOH) intake (g/kg/24 h); (C) preference for EtOH; (D) total fluid intake. Dashed line represents when injections were resumed during the EOD procedure. Data represented as mean ± s.e.m. (**p<0.0021, *p<0.05 for Bonferroni post-hoc tests, n=10 per group).

To determine whether the effect of poly(I:C) on alcohol intake was long lasting, we halted injections after day 29 and measured ethanol intake for another 5 ethanol drinking sessions. During this “no injection” period, there was still a significant main effect of poly(I:C) on ethanol preference [Figure 2C, F_treatment_ (1,18)=6.78, p=0.018; F_time_ (4,72)=4.37, p=0.0032; F_treatment × time_ (4,72)=1.76, p=0.15] and a trend toward increased ethanol consumption [Figure 2B, F_treatment_ (1,18)=3.21, p=0.08; F_time_ (4,72)=0.61, p=0.65; F_treatment × time_ (4,72)=1.286, p=0.28]. Together these results indicate that prior treatment with poly(I:C) can produce a persistent increase in ethanol preference over time.

### 3.3 Poly(I:C) increases alcohol intake across various doses

We investigated whether the effects of poly(I:C) on alcohol intake were dose-dependent. Mice were administered poly(I:C) (2mg/kg or 10mg/kg, i.p.) or saline every four days during EOD consumption of 15% ethanol for 36 days (Figure 3A). Similar to 5mg/kg poly(I:C), multiple injections of these doses were required to increase ethanol consumption [Figure 3B, F_treatment × time_ (16,296)=2.86, p=0.0002]. A similar pattern was observed for ethanol preference [Figure 3C, F_treatment × time_ (16,296)=2.45, p=0.001].

There was no change in total fluid intake [Figure 3D, F_treatment_ (2,37)=0.50, p=0.61; F_time_ (8,296)=9.63, p<0.0001; F_treatment × time_ (16,296)=0.73, p=0.76]. There was also no significant difference between low dose poly(I:C) (2mg/kg) and high dose poly(I:C) (10mg/kg)-treated animals drinking 15% ethanol.

These results suggest that poly(I:C) can increase alcohol intake across the three doses that we tested. However, because ethanol preference was nearly 1.0 when animals were provided 15% ethanol, we switched animals to 20% ethanol during a “no injection” period (beginning at day 38) to reduce ethanol preference and avoid a ceiling effect. During the 16-day “no injection” period the 10mg/kg poly(I:C) group retained increased ethanol consumption compared with saline-treated animals [Figure 3B, F_treatment × time_ (8,148)=2.56, p=0.01]. There was a significant difference in consumption between the 10mg/kg poly(I:C) group and 2mg/kg poly(I:C) group only for the first day of the “no injection” period (p<0.01). Together this suggests that the effects of poly(I:C) on ethanol consumption when poly(I:C) is not repeatedly injected are partially dose-dependent.

We next investigated whether repeated poly(I:C) injections after a period without TLR3 activation would induce similar increases in ethanol drinking. After the “no injection” period, mice were again administered poly(I:C) (2mg/kg or 10mg/kg) or saline every four days during EOD consumption of 20% ethanol for 36 days (beginning at day 59). Both doses of poly(I:C) increased ethanol consumption and preference [Figure 3B ethanol intake: F_treatment × time_(18,333)=2.18, p=0.003); Figure 3C preference: F_treatment × time_(18,333)=1.58, p=0.06]. However, repeated injections of poly(I:C) did not continue to increase alcohol intake *ad infinitum.* Instead, alcohol intake plateaued over time, suggesting that there was an upper limit to how much poly(I:C) can increase drinking.

### 3.4 Poly(I:C) increases alcohol intake only during ethanol drinking

We next investigated whether poly(I:C) alone was sufficient to increase alcohol intake. We administered poly(I:C) (5mg/kg, i.p.) or saline every four days for a total of eight injections before allowing mice to begin 20 days of EOD ethanol (15%) drinking (Figure 4A). Pretreatment with poly(I:C) did not increase ethanol consumption [Figure 4B, F_treatment_ (1,20)=1.584, p=0.22; F_time_ (4,80)=1.024, p=0.40; F_treatment × time_ (4,80)=1.02, p=0.4], although there was a slight initial increase in ethanol preference [Figure 4C, F_treatment × time_(4,80)=3.015, p=0.02], suggesting that poly(I:C) pretreatment changed the pattern of escalation compared with saline-pretreated mice. Total fluid intake was unaffected by poly(I:C) pretreatment [Figure 4D, F_treatment_(1,20)=0.01, p=0.9; F_time_(4,80)=1.01, p=0.42; F_treatment × time_(4,80)=1.65, p=0.17]. Re-administration of poly(I:C) to these animals during access to ethanol increased ethanol consumption [Figure 4B, F_treatment × time_(7,140)=2.7, p=0.01] but did not change ethanol preference [Figure 4C, F_treatment_(1,20)=2.84, p=0.11; F_time_(7,140)=1.233, p=0.29; F_treatment × time_(7,140)=1.38, p=0.22]. Although there was a trend towards reduced total fluid consumption, this effect was not statistically significant [Figure 4D, F_treatment_(1,20)=3.778, p=0.06; F_time_(7,140)=2.55, p=0.01; F_treatment × time_(7, 140)=1.06, p=0.39]. These results suggest that the ability of poly(I:C) to increase alcohol intake depends on the presence of ethanol.

### 3.5 Repeated poly(I:C) during ethanol drinking alters innate immune transcript abundance

Innate immune response transcripts are increased in the brain after chronic exposure to ethanol or poly(I:C) and these increases are thought to drive ethanol consumption (1, 3, 13, 29, 46, 47). To test the hypothesis that poly(I:C)-induced changes in alcohol intake were due to activation of innate immune signaling, we measured changes in the abundance of transcripts for TRIF- and MyD88-dependent genes and for proinflammatory mediators in mice repeatedly administered 5mg/kg poly(I:C) every four days while consuming 15% ethanol in an EOD procedure (Supplemental Figure 5A). The values for each transcript, represented as the fold-change from the respective saline group, are shown as a heatmap in Figure 5A. Raw qRT-PCR data for each transcript are available in Supplemental Figure 5B. Poly(I:C) plus ethanol increased the abundance of transcripts for TRIF-dependent pathway genes (*Tlr3, Ticam1, Ikk*ε*, Irf3*) and proinflammatory mediators (*Ifn*β*1, Ccl5, Ccl2, Il6, Il1*β). In contrast, poly(I:C) plus ethanol decreased the abundance of MyD88-dependent pathway transcripts (*Tlr4, Myd88, Ikk*β). *Tlr2* and *Ifnar1* transcripts were unchanged by treatment.

**Figure 5:**
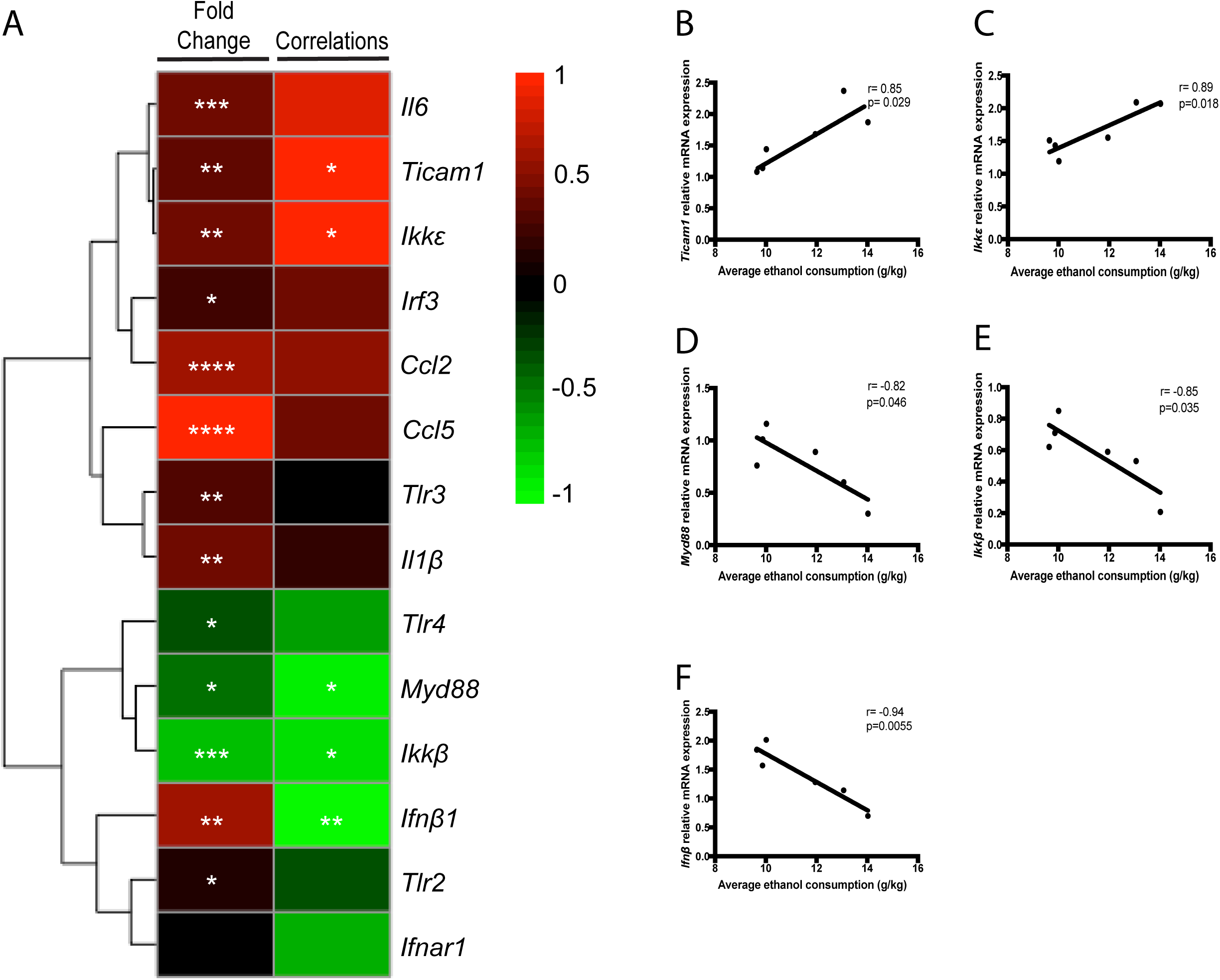
Chronic ethanol and poly(I:C) alter innate immune transcript abundance. Male C57BL6/J mice underwent 36 days of two-bottle-choice, every-other-day drinking (EOD, 15%) with poly(I:C) (5mg/kg) injections every four days for a total of eight injections. Mice were euthanized and prefrontal cortex dissected 24 hours after the final drinking session on day 36. (A) Heat map of innate immune gene transcript abundance and correlations after both poly(I:C) and ethanol. The levels of transcripts are presented as fold-change values (Log2 format) normalized to respective saline-treated groups. The red and green colors indicate high and low transcript abundance or correlation strength. There were significant negative correlations between *Myd88* (B) and amount of ethanol consumed and between *Ikk*β (C) and amount of ethanol consumed in poly(I:C)-treated mice. There were significant positive correlations between *Ticam1* (D) and amount of ethanol consumed and between *Ikk*ε (E) and amount of ethanol consumed in poly(I:C)-treated mice. There was a significant negative correlation between *Ifnb1* (F) and amount of ethanol consumed in poly(I:C)-treated mice. (****p<0.0001, ***p<0.0002, **p<0.0021, *p<0.05 for Tukey post-hoc tests and correlations, n=6 per group).

To identify potential targets that regulate poly(I:C)-induced escalations in drinking, we examined correlation between transcript abundance and average ethanol consumption. Several transcripts were significantly affected by poly(I:C) and alcohol exposure, but only five transcripts correlated with ethanol consumption. In the TRIF-dependent pathway, there was a significant positive correlation (r= 0.85, p=0.029) between *Ticam1* transcript levels and ethanol intake in poly(I:C) treated-mice (Figure 5B). Transcript abundance for the downstream component *Ikk*ε was also positively correlated (r=0.89, p=0.018) with ethanol intake (Figure 5C). Interestingly, there was a negative correlation between Myd88-dependent pathway components and ethanol intake (Figure 5D-E, *Myd88*, r=−0.82, p=0.046; *Ikk*β, r=-0.85, p=0.035). There was also negative correlation (r= −0.94, p=0.0055) between type-1 interferon (*Ifn*β) and alcohol intake. Together these data suggest that both the MyD88- and TRIF-dependent pathways may regulate poly(I:C)-induced escalations in alcohol intake.

### 3.6 Poly(I:C) increases alcohol intake independent of MyD88

Poly(I:C) can act via both MyD88-dependent and –independent mechanisms (42, 48). Moreover, male *Myd88* global knockouts consume more ethanol than littermate controls (18). The strong negative correlation between poly(I:C)-induced escalations in ethanol intake and MyD88-dependent transcript abundance suggested that decreases in MyD88 may be necessary for poly(I:C)-induced increases in alcohol intake. If this were true, then administration of poly(I:C) should not increase alcohol intake in male *Myd88* knockout mice. To test this hypothesis, we administered poly(I:C) (5mg/kg, i.p) every four days to *Myd88* (−/−) knockout mice consuming 15% ethanol in an EOD procedure for 24 days. Poly(I:C) increased ethanol consumption [Figure 6A, F_treatment_ (1,16)=10.99, p=0.0044; F_time_ (5,80)=4.24, p=0.002; F_treatment × time_ (5,80)=1.3, p=0.27] and preference [Figure 6B, F_treatment_(1,16)=14.58, p=0.001; F_time_(5,80)=4.86, p<0.001; F_treatment × time_(5,80)=1.73, p=0.14], but not total fluid intake [Figure 6C, F_treatment × time_=1.97, p=0.09]. These results suggest that poly(I:C) increases alcohol intake independent of MyD88.

**Figure 6:**
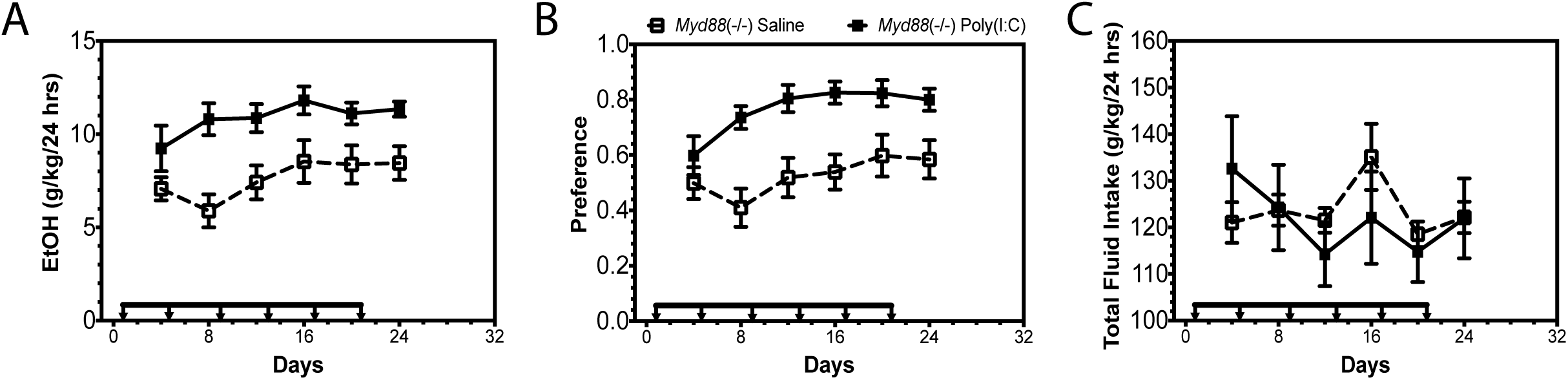
Poly(I:C) increases alcohol intake independent of MyD88. Mutant male mice (*Myd88 −/−*) were injected with saline or poly(I:C) (5mg/kg) every four days for a total of six injections during a two-bottle choice every-other-day drinking procedure (EOD, 15%). (A) Ethanol (EtOH) intake (g/kg/24 h); (B), preference for EtOH; (C) total fluid intake (n=10 per group).

### 3.7 Inhibition of the TLR3-dsRNA complex reduces poly(I:C)-induced increases in alcohol intake

The strong positive correlation between poly(I:C)-induced escalations in alcohol intake and TRIF-dependent transcript abundance suggested that increases in TLR3-dependent signaling may be necessary for poly(I:C)-induced increases in alcohol intake. Additionally, as shown above, poly(I:C)-induced increases in proinflammatory transcripts are partially dependent on TLR3. If TLR3-dependent signaling regulates poly(I:C)-induced drinking behavior, then administration of a TLR3/dsRNA complex inhibitor should decrease alcohol intake in mice chronically administered poly(I:C). To test this hypothesis, we pretreated with saline/poly(I:C) (5mg/kg, i.p) every four days to male C57BL6/J mice consuming 15% ethanol in an EOD procedure for 36 days (Supplemental Figure 6). Once we verified poly(I:C) increased alcohol intake, we injected TLR3/dsRNA complex inhibitor (2mg, i.p.) then injected poly(I:C) (5mg/kg, i.p.) and allowed mice to undergo eight days EOD drinking. Mice treated with a TLR3/dsRNA complex inhibitor before poly(I:C) injection decreased ethanol consumption (Figure 7A). TLR3/dsRNA complex inhibition did not alter preference or total fluid intake in poly(I:C)-treated animals (Figure 7B-C). We further tested the hypothesis that TLR3 inhibition regulates alcohol intake by administering the TLR3/dsRNA inhibitor alone and monitoring drinking behavior. We found that the TLR3/dsRNA inhibitor alone reduced alcohol preference and slowed escalation of alcohol intake (Supplemental Figure 7). Together these results suggest that poly(I:C)-induced increases in alcohol intake are partially dependent on TLR3.

**Figure 7:**
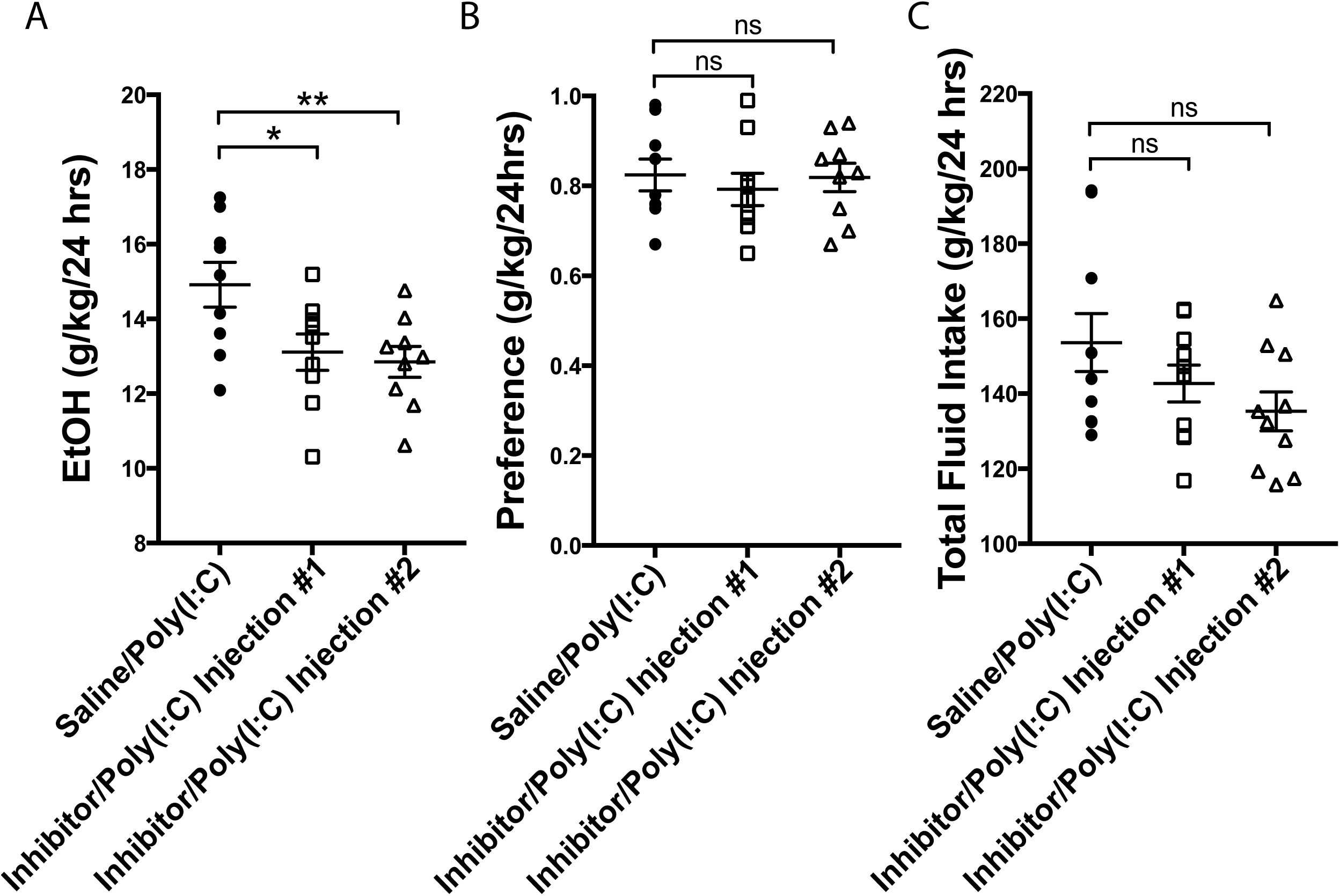
TLR3/dsRNA complex inhibitor reduces poly(I:C)-induced increases alcohol intake. C57BL/6J male mice were injected with saline(10%DMSO) or poly(I:C) (5mg/kg) every four days for a total of nine injections during EOD (15%v/v). Once poly(I:C) increased alcohol intake above saline controls, we injected TLR3/dsRNA complex inhibitor (2mg, i.p.) then injected poly(I:C) (5mg/kg, i.p.) and allowed mice to undergo eight days EOD. (A) Ethanol (EtOH) intake (g/kg/24 h); (B) preference; (C) total fluid intake (g/kg/24h). There was a significant decrease in alcohol intake after each inhibitor pretreatment (Injection #1: t(16)=2.336, p=0.033; Injection #2, t(16)=2.83, p=0.013). Preference (Injection #1: t(16)=0.12, p=0.91; Injection #2, t(16)=0.64, p=0.52) and total fluid intake (Injection #1: t(16)=1.19, p=0.25; Injection #2, t(16)=1.96, p=0.07) were unaffected by treatment. Student’s *t*-tests (*p<0.05, **p<0.02 compared with saline/poly(I:C) group, n=10 per group).

**Figure 8:**
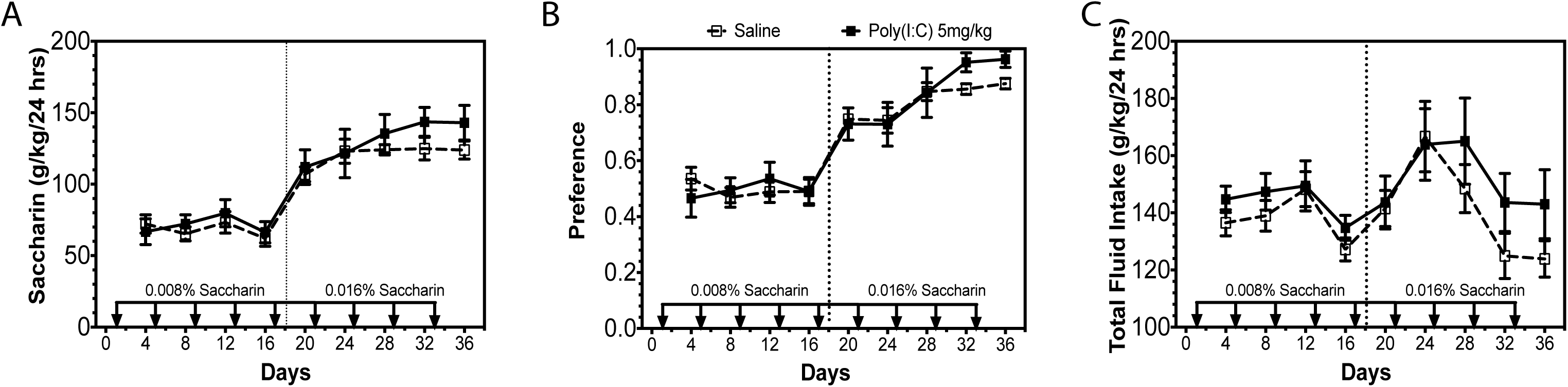
Poly(I:C) does not change taste perception. C57BL/6J male mice were injected with saline or poly(I:C) (5mg/kg) every four days for a total of nine injections during a two-bottle choice every-other-day saccharin procedure (0.0008% or 0.016%). Dashed line represents change in saccharin concentration. (A) Saccharin consumption (g/kg/24 h); (B) preference for saccharin; (C) total fluid intake (g/kg/24 h) (n=10 per group).

### 3.8 Poly(I:C) does not alter saccharin consumption

We also studied consumption of saccharin using an EOD procedure to determine whether altered sweet taste perception could account for changes in ethanol consumption. Poly(I:C) did not change EOD saccharin intake at either 0.0008% [Figure 7A, F_treatment_(1,12)=0.14, p=0.71; F_time_(3,36)=1.81, p=0.16; F_treatment × time_(3,36)=0.58, p=0.63] or 0.016% saccharin [Figure 7A, F_treatment_(1,12)=1.06, p=0.32; F_time_(4,48)=3.07, p=0.02; F_treatment × time_(4,48)=0.58, p=0.68]. Additionally, poly(I:C) did not change saccharin preference at either 0.0008% [Figure 7B, F_treatment_(1,12)=0.0001, p=0.98; F_time_(3,36)=0.24, p=0.86; F_treatment × time_(3,36)=0.91, p=0.44] or 0.0016% saccharin [Figure 7B, F_treatment_(1,12)=0.55, p=0.47; F_time_(4,48)=7.78, p<0.0001; F_treatment × time_(4,48)=0.83, p=0.51]. Total fluid intake was also unchanged by poly(I:C) [Figure 7C, F_treatment_(1,12)=1.14, p=0.31; F_time_(8,96)=5.55, p<0.0001; F_treatment × time_(8,96)=0.68, p=0.70]. These findings indicate that poly(I:C) does not change detection of sweet taste, suggesting that poly(I:C) does not increase alcohol intake by perturbing saccharin taste perception.

## 4. Discussion

Our results reveal a novel role for TLR3-dependent signaling in regulation of alcohol intake. Activation of TLR3 by poly(I:C) increased transcript abundance of innate immune signaling components in prefrontal cortex. Repeated TLR3 activation, across multiple doses, increased alcohol intake over time. However, the effect of TLR3 activation on drinking was dependent on both repeated injections and the presence of ethanol, suggesting poly(I:C) is necessary for escalations in alcohol intake but not sufficient when administered alone. Poly(I:C) increased alcohol intake without altering saccharin taste preference, suggesting changes in consumption were not due to alterations in sweet taste preference. Ethanol and poly(I:C) exposure increased transcript abundance of both TRIF- and MyD88-dependent pathway components. TRIF-dependent pathway levels positively correlated with increased alcohol intake and MyD88-dependent pathway levels negatively correlated with alcohol intake. Poly(I:C) escalated alcohol intake in *Myd88* knockout mice, supporting the hypothesis that increases in alcohol intake after poly(I:C) occur independently of MyD88. Inhibition of the TLR3/dsRNA complex reduced escalation of alcohol intake and poly(I:C)-induced escalations in alcohol intake, supporting the conclusion that increases in alcohol intake after poly(I:C) are partially dependent on TLR3/TRIF-dependent signaling. Together, these results suggest that activation of TLR3 signaling promotes alcohol consumption and therefore inhibition of TLR3 may be a potential therapeutic target for treating excessive drinking.

Poly(I:C) produced transient activation of proinflammatory mediators in prefrontal cortex, with peak cytokine activation occurring at 3 hours post-injection—consistent with previous reports (27, 29, 49). However, poly(I:C) also increased transcript abundance of both TRIF- and MyD88-dependent components across a 48-hour period, suggesting that these longer lasting changes in innate immune signaling may drive drinking behavior. These poly(I:C)-induced changes in innate immune signaling were, at least partially, dependent on TLR3, because inhibition of the TLR3/dsRNA complex reduced activation of proinflammatory mediators. Staining for poly(I:C) presence in the liver and prefrontal cortex confirmed poly(I:C) can enter brain, which is consistent with poly(I:C) being able to disrupt the blood-brain barrier to permit direct brain access for dsRNA (50). This suggested that direct activation of TLR3 in the central nervous system contributes, at least partially, to the behavioral effects of poly(I:C). However, the greater amount of poly(I:C) signal detected in the liver raised the possibility that peripheral actions of poly(I:C) may play a role in how the innate immune system regulates drinking behavior and ethanol metabolism. Viral infections do alter liver metabolism (51). Activation of TLR3 by dsRNA induces natural killer cell accumulation and activation in the liver, leading to liver inflammation and injury (51). However, it has been shown that TLR3 activation with poly(I:C) ameliorates alcoholic liver injury via the stimulation of IL-10 production in hepatic stellate cells and Kupffer cells (52). Moreover, poly(I:C) does not alter alcohol dehydrogenase (ADH) expression in liver (52). Additionally, TLR3 knockout mice show no differences in ethanol metabolism when compared with littermate controls (data not shown). Therefore, if the absence of TLR3 does not affect ethanol metabolism, poly(I:C) does not effect ADH expression in liver and poly(I:C) treatment ameliorates alcoholic liver injury, we hypothesize that poly(I:C) is not changing ethanol metabolism. However, future studies will need to dissect the role of peripheral versus central TLR3 in the regulation of alcohol intake.

Repeated injections of poly(I:C) were required to increase alcohol intake, suggesting that long-lasting alterations in gene expression must occur to change behavior rather than transient innate immune changes due to acute TLR3 activation. All tested doses increased alcohol intake over time, indicating no dose-dependence of poly(I:C) on alcohol intake. For both the 5mg/kg and 10mg/kg doses, poly(I:C) elevated alcohol preference even when poly(I:C) injections were halted, consistent with a persistent change in innate immune signaling having occurred. However, alcohol intake did not continue to increase without continued poly(I:C) injections, indicating that poly(I:C) must be repeatedly injected to increase alcohol consumption. The requirement of repeated TLR3 activation suggests different and longer lasting signaling changes must occur to increase alcohol intake. A limitation of the current study is that a single time point was selected after poly(I:C) and ethanol consumption (24h), which provided a snapshot of how poly(I:C) changes gene expression. From these data we hypothesize that TRIF-dependent signaling may be necessary for escalations in alcohol intake along with longer-lasting changes in innate immune signaling pathways not investigated in this paper. Future studies will investigate the longer-lasting transcriptional changes necessary for poly(I:C)-induced increases in alcohol intake over time using RNA-sequencing in order to address how repeated TLR3 activation changes drinking behavior on the transcriptome level.

To determine if poly(I:C) alone is sufficient to increase alcohol intake we pretreated animals with poly(I:C) for eight injections before allowing animals to begin drinking. Pretreatment with poly(I:C) did not change alcohol intake, suggesting that poly(I:C) alone is not sufficient to change drinking behavior. Additionally after pretreatment, we demonstrate ethanol and poly(I:C) must be present together to increase alcohol intake, implying that ethanol adds a key component of neuroimmune activation that we have yet to discover. It has been hypothesized that alcohol-induced neuroinflammation directly contributes to the development of alcohol use disorders (53). Ethanol can activate TRIF-dependent and MyD88-dependent signaling as well as a wide array of proinflammatory mediators (2, 54). Others have shown that a variety of proinflammatory signals increases ethanol drinking and preference (13) and interfering with neuroinflammatory signaling reduces alcohol consumption (55, 56). Combined activation of TLRs alters the extent of cytokine production and can stimulate additive increases in the amount of cytokines produced, which could lead to greater increases in alcohol consumption over time (57–59). Thus, the combination of ethanol and poly(I:C) may be necessary to produce sufficient inflammatory signaling to change drinking behavior.

Repeated ethanol and poly(I:C) exposure altered innate immune transcript abundance in both TLR pathway branches. Additionally there was a robust increase in levels of proinflammatory mediators (*Il6, Il1*β*, Ccl2, Ccl5)* in poly(I:C)-treated mice, suggesting involvement of multiple innate immune signaling branches. When there is chronic activation of one immune signaling branch (e.g. TRIF-dependent), there can be compensatory downregulation of the other (e.g. MyD88-dependent) (60). This is thought to allow the innate immune system to facilitate simultaneous amplification of the appropriate level of protective responses during infection, attenuation of the damage inflicted by inflammation, and maintenance of homeostasis (61, 62). For example, poly(I:C) can downregulate TLR4 signaling via a TLR3-dependent mechanism to attenuate the immune response (63, 64). We considered whether TLR3 induced decreases in MyD88-dependent signaling could explain the increase in alcohol intake that we observed in mice treated with poly(I:C). In support of this hypothesis, knockout of *Myd88* increases alcohol intake in male mice (20, 21). However, we found that poly(I:C) increased alcohol intake in *Myd88* knockout mice—a limitation of this knockout study was that wildtype controls were not used in conjuction with *Myd88* knockout mice. Taken together, this suggests that poly(I:C) increases alcohol intake through mechanisms independent of MyD88.

TLR3 activation can have either neuroprotective or neurodegenerative effects. For instance, poly(I:C) pretreatment exerts neuroprotective and anti-inflammatory effects in simulated cerebral ischemia models (65, 66). However, in a Parkinson’s model, poly(I:C) triggered nigrostriatal dopaminergic degeneration and microglial proliferation (67, 68). This suggests that when inflammation has already occured, TLR3 activation primes immune cells in brain for proinflammatory responses leading to enhanced neurodegeneration. We have previously shown EOD drinking enhances proinflammatory signaling (24), thus we hypothesized that poly(I:C) increases alcohol intake by enhancing inflammatory responses, specifically TLR3/TRIF-dependent responses. In support of this hypothesis we observed increases in TRIF-dependent pathway components after chronic poly(I:C) and ethanol, which positively correlated with alcohol intake. Furthermore, inhibition of the TLR3/dsRNA complex reduced drinking in poly(I:C)-treated animals, indicating that TLR3/TRIF-dependent signaling may promote excessive alcohol intake but other immune pathways may also mediate this effect. Global knockout of TLR3 reduces EOD drinking in male mice (Blednov unpublished observations) and pharmacological inhibition of the downstream TRIF-dependent pathway components IKKI and TBK1 decreases alcohol intake (24). TLR3 signaling may also have broader roles in regulating rewarding effects of drugs. For instance, TLR3 has been shown to modulate the behavioral effects of cocaine in mice and intra-nucleus accumbens injection of TLR3/dsRNA complex inhibitor significantly attenuated cocaine self-administration in mice (69). TLR3 activation also enhances morphine conditioned place preference acquisition (70). Therefore, poly(I:C) may increase alcohol intake by changing the rewarding properties of ethanol. Future studies will need to dissect if TLR3 activation changes ethanol sensitivity. Taken together, inhibition of TLR3 may be a valuable therapeutic target to reduce excessive alcohol consumption as well as treating other substance abuse disorders.

Although poly(I:C) is a synthetic dsRNA polymer that mimics viral infection, it should be noted that we are most likely not modeling viral infection in human alcoholics. In human alcoholics, chronic alcohol exposure over time interferes with the normal functioning of all aspects of the innate and adaptive immune response, including both cell-mediated and humoral responses (71). Most likely our model represents a heightened peripheral proinflammatory state, driven by TLR3 that leads to increases in alcohol intake. Since poly(I:C) injection results in more dsRNA signal in the liver compared with prefrontal cortex, we are likely activating more peripheral TLR3 resulting in an increase in proinflammatory mediators in the periphery that can cross the BBB and activate central neuroimmune pathways (such as IL-1β, IL-6, and IFNβ).

Proinflammatory cytokines have been detected in the serum of human alcoholics and higher levels of cytokines correlates with alcohol craving (11). Therefore, we hypothesize that these proinflammatory mediators increase craving and increase alcohol intake over time. Future studies should address whether heightened viral load increases alcohol consumption in human populations, linking the idea that viral activation of TLR3 leads to escalation of alcohol intake over time.

## 5. Conclusions

Altogether, this work has defined the role of TLR3 signaling in drinking behavior, defined the causal role and pathways necessary for TLR3-dependent increases in alcohol intake, and provides a potential pharmacotherapeutic target to improve treatment outcomes for excessive drinking. While, these studies suggest that increases in TLR3-dependent signaling exacerbate proinflammatory signaling resulting in increases in alcohol intake, we suggest targeted inhibition of TLR3 and its downstream mediators may represent novel treatment strategies for excessive drinking.

## Supporting information

## Funding

This work was supported by the National Institutes of Health/National Institute of Alcohol Abuse and Alcoholism [U01 AA020926, P01 AA020683, AA013520, AA006399, AA025499]. The authors report no biomedical financial interests or potential conflicts of interest.

---

Supplemental Figure 1: *K1 antibody validation*. (A) Staining specificity for K1 antibody in prefrontal cortex and liver of C57BL/6J male mice injected with R848 (ssRNA TLR7/8 agonist). (B) Secondary controls in prefrontal cortex and liver of C57BL/6J male mice— primary antibody omitted, 10% donkey serum and donkey anti-mouse Alexa Flour 488 secondary. (C) Labeling staining controls in prefrontal cortex and liver of C57BL/6J male mice—primary antibody and secondary antibody omitted, 10% donkey serum only. Scale bar=50μm (n=4 per group).

Supplemental Figure 2: *Raw qRT-PCR data for poly(I:C) timecourse*. The levels of transcripts are presented using fold-change values normalized to endogenous control, with six biological and three technical replicates. Data represented as fold change ± s.e.m (****p<0.0001, ***p<0.0002, **p<0.0021, *p<0.05 compared with saline treatment at same time point; ^#^p<0.05 compared with other poly(I:C) groups for Tukey post-hoc tests, n=6 per group).

Supplemental Figure 3: *TLR3/dsRNA complex inhibitor reduces poly(I:C)-induced increases in proinflammatory transcript abundance*. The levels of transcripts are presented using fold-change values normalized to endogenous control, with six biological and three technical replicates. Values are presented as fold change ± SEM. (* represents significant difference from saline group, # represents significant difference from poly(I:C) group, ****p<0.0001, ***p<0.0002, **p<0.0021, *p<0.05 compared with saline treatment at same time; ^#^p<0.05 compared with other poly(I:C) groups, n=6 per group).

Supplemental Figure 4: *dsRNA-like immunoreactivity is detectable in both prefrontal cortex and liver after poly(I:C) administration*. C57BL/6J male mice were injected with poly(I:C) (5mg/kg). Mice were euthanized, prefrontal cortex dissected and liver removed three hours post-injection. (A-D) representative images of K1 dsRNA immunofluorescence, scale bars=100μm. (E) Corrected total cell fluorescence (CTCF) was increased after poly(I:C) injection in both brain and liver, though the increase was greater in the liver [F_treatment_(1,12)=665, p<0.0001; F_organ_(1,12)=287, p<0.0001; F_treatment × organ_(1,12)=327, p<0.0001]. (F) As similar pattern was observed for measurements of integrated intensity [F_treatment_(1,12)=80.45, p<0.0001; F_organ_(1,12)=122,4, p<0.0001; F_treatment × organ_(1,12)=31.91, p<0.0001]. (G) Poly(I:C) treatment increased the number of dsRNA immunopositive cells present in the prefrontal cortex (two-tailed Student’s *t*-test t(6)=25.73, ****p<0.0001). All data are expressed as mean ± s.e.m, n=4 per group.

Supplemental Figure 5: *Poly(I:C) and ethanol increase alcohol intake and change innate immune transcript abundance* (A) Drinking data for poly(I:C) (5mg/kg) EOD experiment used for qRT-PCR analysis. There was a main effect of poly(I:C) on ethanol intake [F_treatment_(1,18)=15.49, p=0.001; F_time_(7,126)=4.65, p=0.0001; F_treatment × time_(7,126)=1.433, p=0.19] and preference [F_treatment_(1,18)=13.54, p=0.001; F_time_(7,126)=5.09, p<0.0001; F_treatment × time_(7,126)=0.37, p=0.92]. There was no effect of poly(I:C) on total fluid intake [F_treatment_(1,18)=0.55, p=0.46; F_time_(7,126)=2.445, p=0.02; F_treatment × time_(7,126)=1.84, p=0.08]. Raw qRT-PCR data for 5mg/kg poly(I:C) EOD experiment. The levels of genes are presented as fold change ± s.e.m. All values were normalized to endogenous control. Data were analyzed by two-tailed, Student’s *t*-tests (****p<0.0001, ***p<0.0002, **p<0.0021, *p<0.05 compared with saline group, n=7 per group).

Supplemental Figure 6: *TLR3/dsRNA complex inhibitor/poly(I:C) treatment drinking data*. C57BL/6J male mice were injected with saline (10%DMSO)/saline or saline(10%DMSO)/ poly(I:C) (5mg/kg) every four days for a total of nine injections during two-bottle choice, every-other-day drinking (EOD,15%), then poly(I:C) treated animals were pretreated with a TLR3/dsRNA complex inhibitor for eight days. (A) Ethanol (EtOH) intake (g/kg/24 h); (B) preference for EtOH; (C) total fluid intake (g/kg/24 h); (D-E) body weight (g). There was a main effect of poly(I:C) on ethanol [F_treatment_(1,17)=7.5, p=0.014; F_time_(8,136)=15.01, p<0.0001; F_treatment × time_(8,136)=2.29, p=0.02] and preference [F_treatment_(1,17)=3.626, p=0.07; F_time_(8,136)=15.89, p<0.0001; F_treatment × time_(8,136)=2.83, p=0.019]. Total fluid intake was not changed by poly(I:C) treatment [F_treatment_(1,17)=0.14, p=0.71; F_time_(8,136)=3.65, p=0.0007; F_treatment × time-_ (8,136)=1.59, p=0.13]. There was a main effect of TLR3/dsRNA inhibitor pretreatment on poly(I:C)-induced ethanol consumption (F_treatment_(1,16)=7.25, p=0.01). There was no effect of TLR3/dsRNA inhibitor pretreatment on poly(I:C)-induced ethanol preference (F_treatment_(1,16)=0.16, p=0.69) or total fluid intake (F_treatment_(1,16)=3.35, p=0.08). There was no main effect of poly(I:C) or TLR3/dsRNA inhibitor on body weight [F_treatment_(1,18)=1.49, p=0.24; F_time_(10,180)=9.4, p<0.0001; F_treatment × time_(10,180)=2.69, p=0.004]. Arrows indicate days when animals received injections, the red arrow indicates switch to the TLR3/dsRNA inhibitor. Dashed line represents when saline(10%DMSO)/poly(I:C) treated animals were treated with inhibitor(10%DMSO)/poly(I:C). Data represented as mean + s.e.m. (*p<0.05, significantly different from saline/saline group; #p<0.05, significantly different from saline/poly(I:C) group, Bonferroni post-hoc tests, n=10 per group).

Supplemental Figure 7: *TLR3/dsRNA inhibitor alone slows escalation of alcohol preference.* Male C57BL6/J mice were treated with either saline(10%DMSO) or TLR3/dsRNA complex inhibitor (2mg) and underwent 24 days of EOD-2BC (15%v/v ethanol). To determine how the inhibitor changed escalation of drinking behavior from baseline we subtracted subsequent behavioral drinking days from the first drinking day. (A) Inhibitor treatment trended toward decreased ethanol intake (F_treatment_(1,18)=3.49, p=0.07; F_time_(4,72)=5.57, p=0.0006; F_treatment × time_(4,72)=0.363, p=0.83). (B) Inhibitor treatment reduced escalation of alcohol preference (F_treatment_(1,18)=5.96, p=0.02; F_time_(4,72)=6.79, p=0.0001; F_treatment × time_(4,72)=1.149, p=0.34). (C) Inhibitor treatment modestly increased total fluid intake escalation (F_treatment_(1,18)=4.92, p=0.04; F_time_(4,72)=3.38, p=0.01; F_treatment × time_(4,72)=2.62, p=0.04). Data represented as mean ± s.e.m. (*p<0.05, significantly different from saline group, Bonferroni post-hoc tests, n=10 per group).

