## Supplementary figures and images for "Toll-like receptor 3 activation increases voluntary alcohol intake in C57BL/6J male mice"

### Supplementary file 1

Warden, A. TLR3 activation increases alcohol intake. Supplemental Figure 1

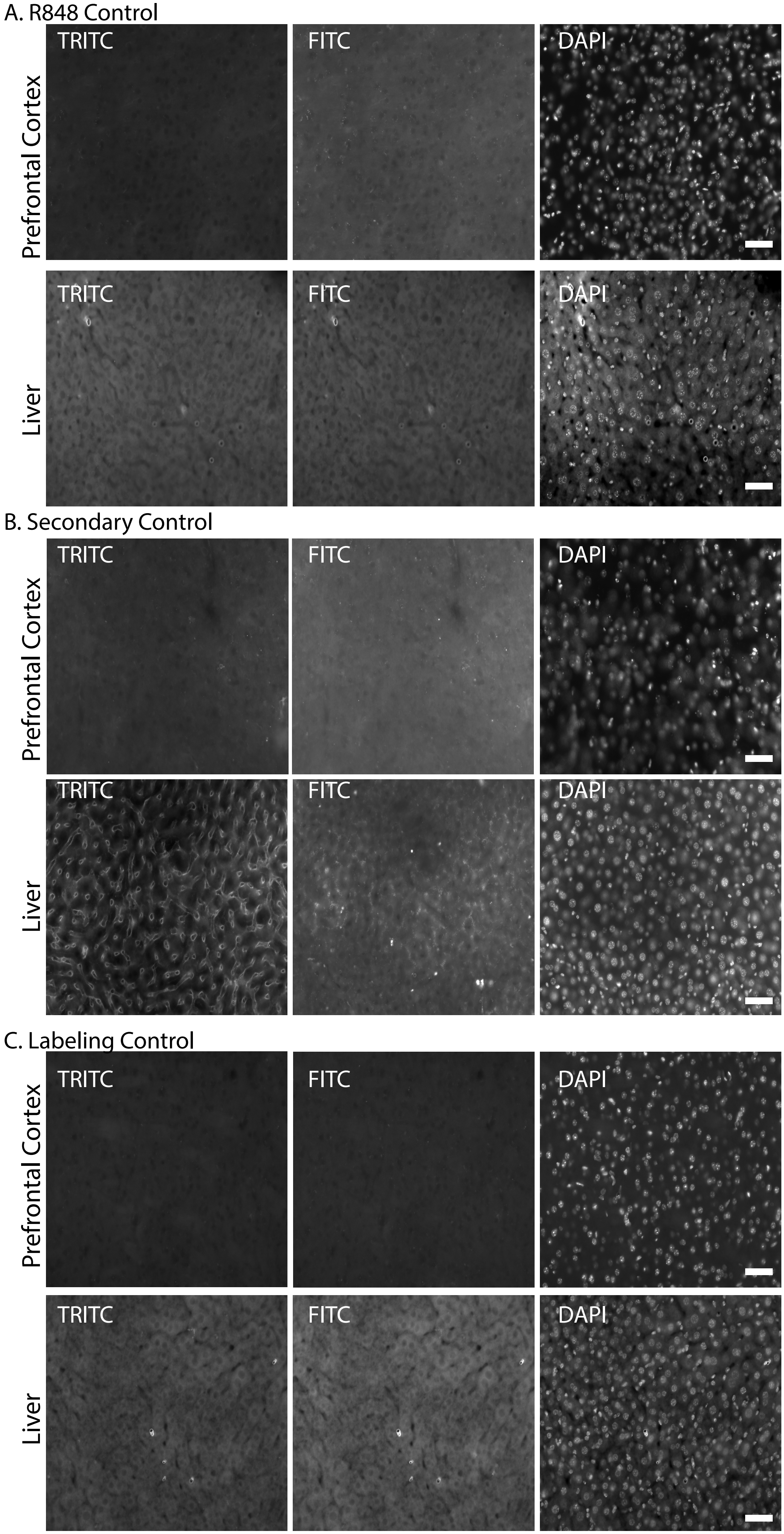

### Supplementary file 2

Warden, A. TLR3 activation increases alcohol intake. Supplemental Figure 2

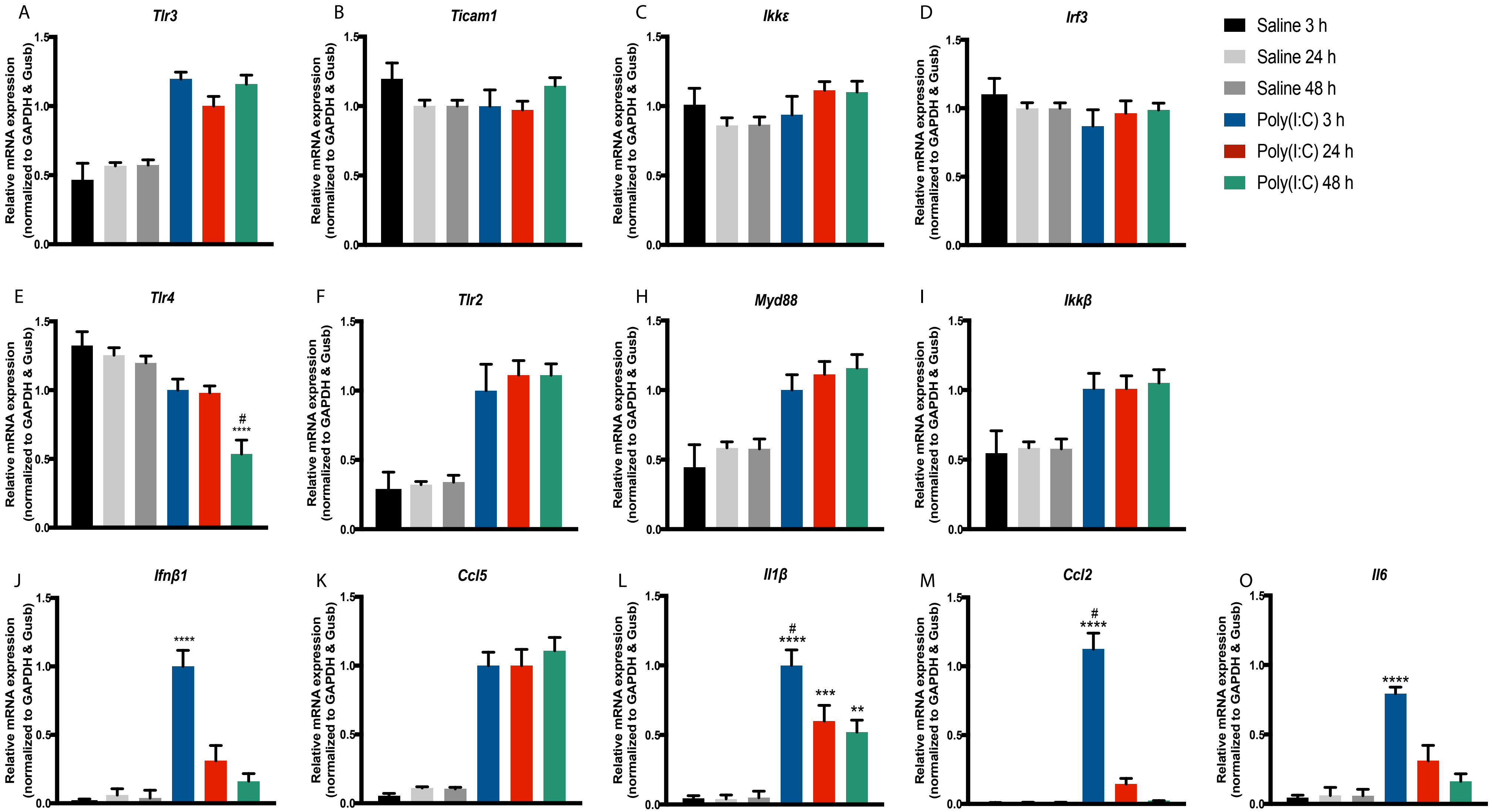

### Supplementary file 3

Warden, A. TLR3 activation increases alcohol intake
Supplemental Figure 3

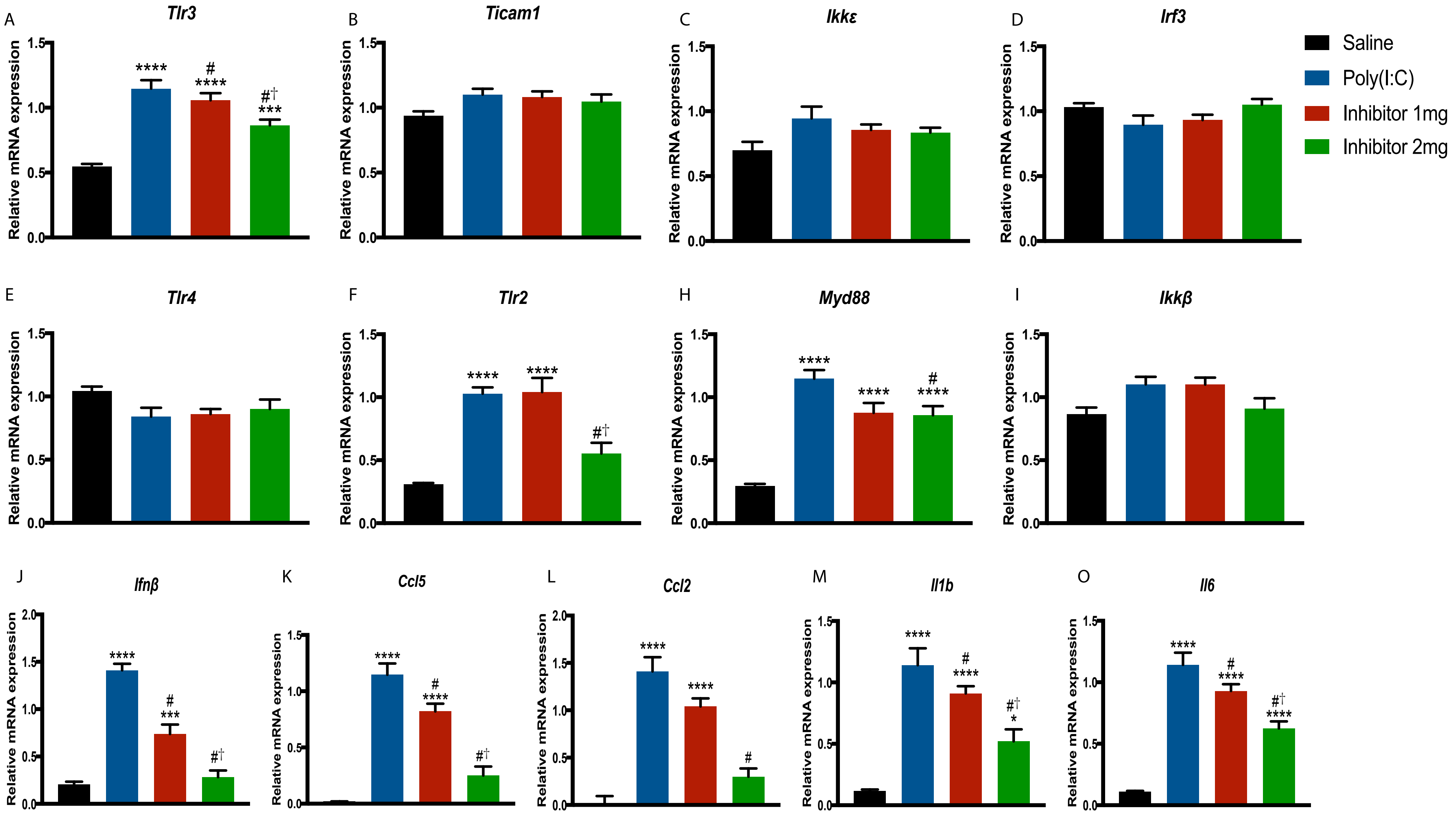

### Supplementary file 4

Warden, A. TLR3 activation increases alcohol intake.
Supplemental Figure 4

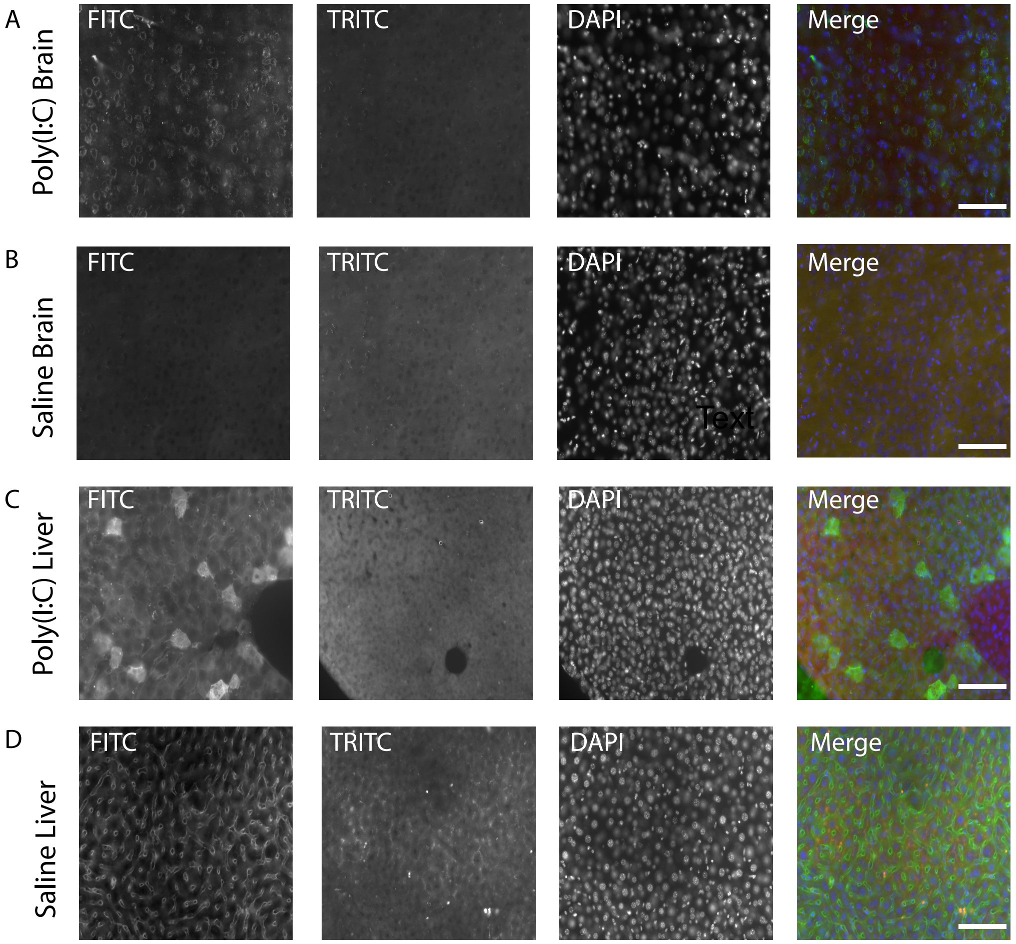


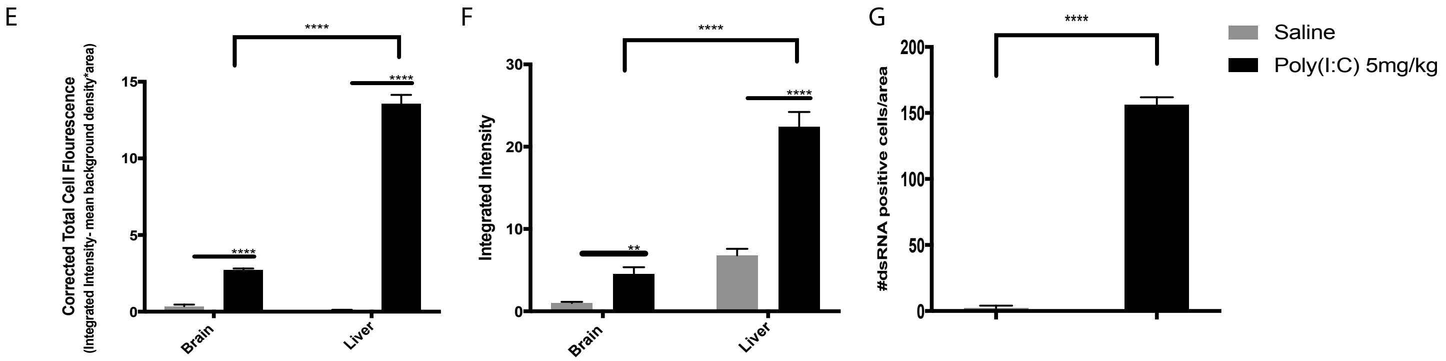

### Supplementary file 5

Warden, A. TLR3 activation increases alcohol intake.
Supplemental Figure 5

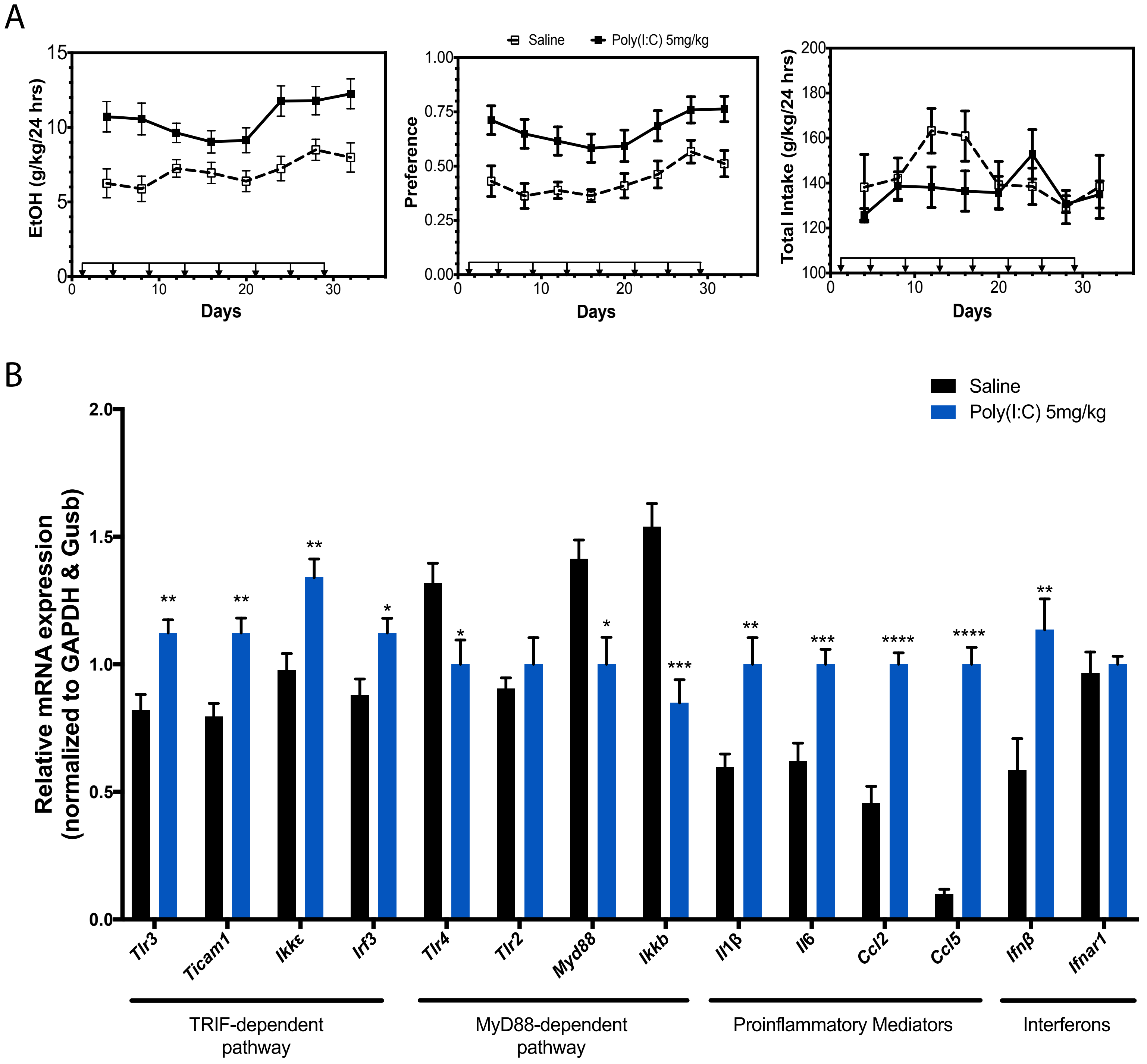

### Supplementary file 6

Warden, A. TLR3 activation increases alcohol intake.
Supplemental Figure 6

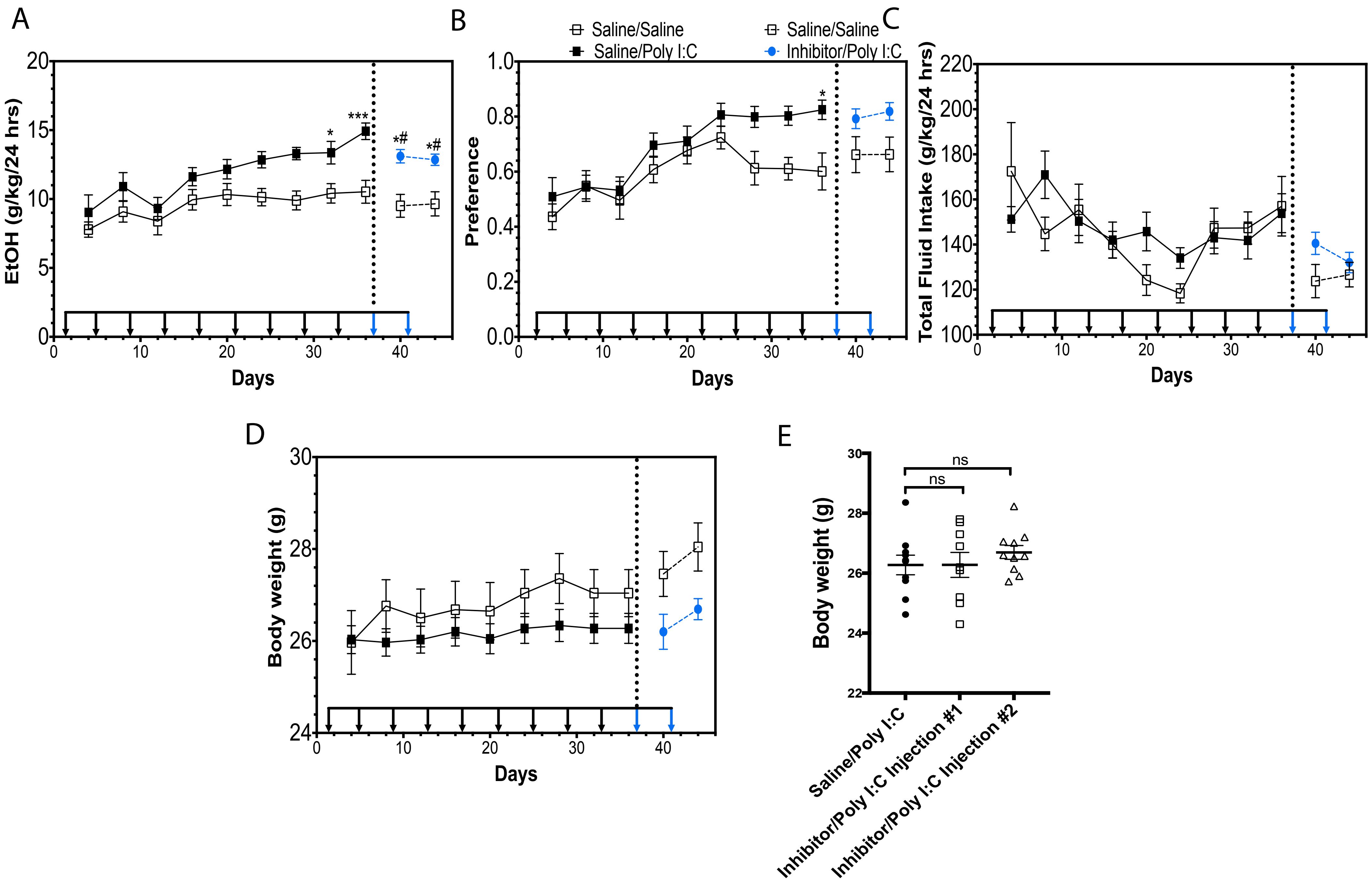

### Supplementary file 7

Warden, A. TLR3 activation
increases alcohol intake.
Supplemental Figure 7

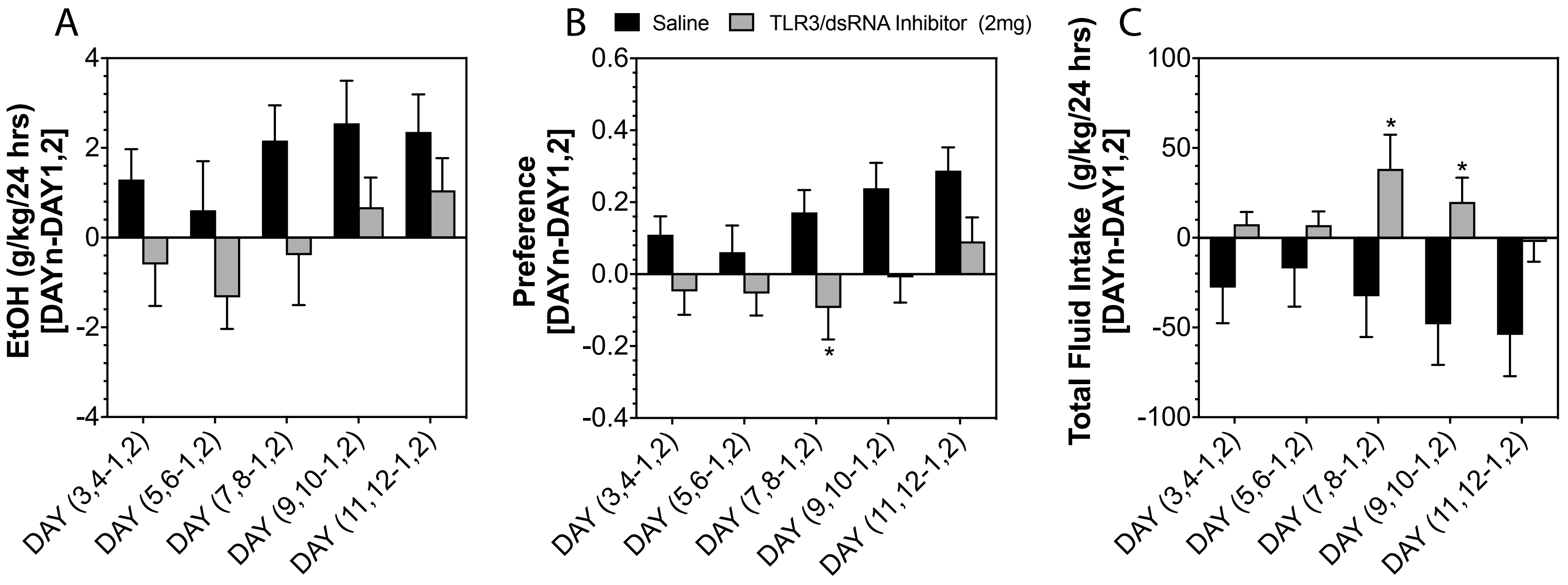
